# Increased circulating TREM2^+^ microglial extracellular vesicles in aged APP/PS1 Alzheimer’s disease rats

**DOI:** 10.1101/2025.06.02.657443

**Authors:** Sarah J. Myers, Brian L. Allman, Stephen H. Pasternak, Shawn N. Whitehead, Austyn D. Roseborough

## Abstract

**INTRODUCTION:** TREM2 is a microglial marker important in Alzheimer’s disease (AD) pathogenesis, but current methods to detect microglial TREM2 expression *in vivo* are limited. Circulating microglia-derived extracellular vesicles (EVs) show promise as potential biomarkers for AD and may offer insight into TREM2 activity.

**METHODS:** TMEM119^+^/TREM2^+^ EVs were assessed using nanoscale flow cytometry in plasma from wildtype and APP/PS1 rats aged to 3-, 9-, and 15-months-old. Molecular and histological assays were used to assess microglia markers in rat brain tissue and a radial arm water maze task was employed to evaluate spatial working and reference memory.

**RESULTS:** Circulating TMEM119^+^/TREM2^+^ EVs were increased in 15-month APP/PS1 rats and associated with severity of cognitive impairment. TREM2 brain expression varied by anatomical region, age, transgene, and assay.

**DISCUSSION:** Collectively, this study provides the first assessment of TMEM119^+^/TREM2^+^ EVs as a biomarker of brain microglial expression and cognition in an AD rat model.

## 1. Introduction

Alzheimer’s disease (AD) is the most common cause of dementia, accounting for at least 60% of the 55 million people living with dementia worldwide^1^. Without curative treatments, the number of people living with dementia globally is predicted to rise to over 150 million by 2050^1^. AD is diagnosed by post-mortem identification of extracellular amyloid beta (Aβ) deposits and intracellular tau neurofibrillary tangles. However, these are not the only pathologies to appear in the AD brain and Aβ plaques are common in individuals without dementia^2,3^, emphasizing the need for further investigation into additional pathologies. For example, recent research has focused on the potential role of microglia, a class of brain cells located throughout the central nervous system (CNS) that work to maintain brain homeostasis through functions such as phagocytosis, synaptic pruning, myelin maintenance, and inflammatory processes. Importantly, both experimental and clinical studies have identified a shift in microglia function as a prominent feature of AD and suggest it begins early in disease progression^4–7^. Multiple roles have been proposed for microglia in the pathogenesis of AD, in which they may be protective through Aβ clearance early in disease, but a transition to a chronic inflammatory state in later disease can be toxic to surrounding neurons^8,9^.

Transcriptomic analyses of human^10–13^ and rodent ^14–16^ brain tissue have identified a diverse range of microglial phenotypes that differ by age, brain region, and disease pathology. Notably, these phenotypes include white matter aging microglia (WAM) and disease-associated microglia (DAM), which are both upregulated in experimental models of AD and are dependent on triggering receptor expressed on myeloid cells 2 (TREM2) signaling^15,17^. TREM2 is a cell surface receptor that functions via its interaction with the DAP12 protein to initiate pathways that promote phagocytosis, survival, chemotaxis, inflammatory responses, and proliferation^17–21^.

Loss-of-function mutations in TREM2 are linked to an increased risk for developing AD, implying that a change in TREM2 function contributes to disease pathogenesis^22,23^. Experimental AD models also suggest an important role for TREM2, as studies show that TREM2 signalling is required for microglial recruitment to amyloid plaques^24,25^. However, the effect of TREM2 deficiency on AD pathology is inconsistent across studies, varying by brain region, age, and model^24–28^. Further investigation into the relationship between TREM2 signalling and AD pathology is required but is challenged by current limitations in methodology for detecting microglial phenotypes *in vivo*.

Aside from post-mortem brain analyses, positron emission tomography (PET) is the only technique available to assess microglial expression *in vivo*; however, current tracers lack cell specificity and the ability to detect specific phenotypes^29^. A promising complementary technique involves the sampling of extracellular vesicles (EVs), which are 100-1000 nm lipid-bound particles released from cells, and have emerged as potential biomarkers for AD and other neurological diseases^30–32^. EVs are important in cell-to-cell signaling, through processes such as cargo transport and presentation of surface markers to target cells. They can also cross the blood-brain barrier carrying the unique proteins, lipids, RNA, and DNA from their cells-of-origin, providing a snapshot into the physiological state of the cell. Past studies evaluating brain-derived EVs in clinical AD have focused mostly on neuron-^33,34^ and astrocyte-derived EVs^35,36^, as well as markers involved in AD pathology such as Aβ and tau isoforms^37^. At present, however, circulating microglia-derived extracellular vesicles (MEVs) in AD have not yet been well described.

TREM2^+^ microglia are a prominent feature of AD that currently cannot be quantified clinically. Thus, our first objective was to identify TREM2^+^ MEVs in wildtype and double-transgenic APP/PS1 rats across age using nanoscale flow cytometry, a high-throughput approach to EV analysis optimized for particles 80-1000 nm in size. Our second objective was to characterize TREM2 expression in the APP/PS1 model, focusing on regions previously shown to have high DAM and WAM expression (hippocampus and corpus callosum) via protein, RNA, and histological brain tissue analyses. Lastly, as previous experimental and clinical studies have identified a relationship between microglial activity and cognition^6,38^, our final objective was to characterize spatial working and reference memory and assess its relationship with TMEM119^+^/TREM2^+^ EVs. Collectively, our results show for the first time that TMEM119^+^/TREM2^+^ EVs can be detected in the systemic circulation, are upregulated in 15-month APP/PS1 rats, and associate with the degree of cognitive impairment.

## 2. Methods

### 2.1 Animals

Animal ethics and procedures used in this study were approved by the Animal Care Committee at Western University (protocol 2022-137). All rats included in this study were housed in facilities maintained by Western University Animal Care and Veterinary Services on a 12-hour light/dark cycle with *ad libitum* access to food and water. Male and female wildtype Fischer 344 and Fischer 344-APP/PS1 double transgenic rats (APP/PS1) were bred and aged in-house to 3-, 9-, or 15-months of age. APP/PS1 rats are homozygous for the human amyloid precursor protein (APP) transgene with Swedish and Indiana mutations and hemizygous for the human presinilin1 (PS1) transgene with an L166P mutation^39^.

### 2.2 Radial arm water maze

#### Setup

We employed a 4/8 radial arm water maze (RAWM) task adapted from Hyde et al.^40^ to assess spatial working and reference memory. In a dimly lit room, a circular tank (144 cm) was filled with room temperature water (22-23 °C) and dyed black with non-toxic acrylic paint. An 8-arm steel maze inset (height: 50.8 cm; arm length: 63.5 cm measured from centre of maze; arm width: 15.24 cm) was placed inside of the circular tank (Maze Engineers). The protocol occurred over 17 consecutive days, where rats underwent 1 day of habituation, 1 day of training, and 15 days of testing. Rats were tested at the same time each day and brought into the room to acclimate ∼30 mins before beginning.

#### Habituation

On the first day, the rats were habituated to the maze over six trials (∼30 s intertrial intervals) which trained them to navigate from the start arm to hidden escape platforms (3 cm below water surface) located at the ends of two of the arms. The tank was enclosed by curtains without the presence of spatial cues. For all six trials, rats swam to locate a platform from various locations and remained on the platform for 30 s before removal from the tank. For trials 1 and 2, the rats were placed in an enclosed maze arm, opposite the hidden platform. Trials 3 and 4 progressed to the rats being placed in the centre of the maze with all but one arm closed off. Finally, rats were placed facing the back of the start arm for trials 5 and 6 and allowed to swim to the open arm with a hidden platform. If rats failed to find the platform within 60 s, they were cued to the platform location.

#### Training

The next day, four visual cues were placed in a square around the maze (yellow cross, white star, orange rectangle, and green triangle) and all maze arms were opened. A given animal had four platforms to find over four training trials (∼30 s intertrial intervals). A trial consisted of the animal being placed facing the back of the start arm, swimming to find a platform and remaining on the platform for 30 s to observe the surrounding visual cues. The arm of the located platform was closed off for the next trial, and this was repeated until all four platforms were located. If rats failed to find a platform within 120 s they were guided to the nearest available platform.

#### Testing

Following training, rats were given four testing trials per day for fifteen consecutive days with the same platform locations. Testing followed the same protocol as training except located platforms were removed from the tank rather than blocked off. For rats to perform well at this task, they needed to 1) avoid entering arms that never contained platforms (reference memory errors), 2) use a win-shift strategy to avoid re-entering arms that previously contained platforms within that session (working memory correct errors), and 3) avoid repeat entries into arms that never contained a platform (working memory incorrect errors). All trials were recorded using ANYmaze tracking software with a webcam (C930e; Logitech) mounted on the ceiling above the water tank. Arm entries were registered as the midpoint of a rat crossing into a given arm and were scored as errors based on the requirements above, with working memory correct and incorrect errors collapsed together for a single metric.

### 2.3 Blood and brain collection

#### Blood collection

Wildtype and APP/PS1 rats were euthanized at 3-, 9-, or 15-months of age via an intraperitoneal injection of pentobarbital (Euthanyl, Bimeda Animal Health Inc). Terminal blood was collected from the left ventricle using an 18-guage needle and deposited into lithium heparin coated Microvette 500 tubes (Sarstedt) to prevent coagulation. Plasma was isolated via two rounds of centrifugation (2500 x *g* for 15 mins at 4°C) and stored at -80 °C. In accordance with the minimal information for studies of extracellular vesicles (MISEV) framework^41^, detailed description of plasma EV handling is provided in sections 2.5 and 2.6.

#### Perfusion

After blood collection, rats underwent transcardial perfusion with 180 mL 0.01 M PBS followed by 300 mL 4% paraformaldehyde (PFA). Brains were collected and stored in 4% PFA for 24 hours at 4 °C and then transferred to 30% sucrose for at least 36 hours prior to sectioning. Brains were cut using a cryostat (CryoStar NX50, Thermo Fisher Scientific) into 30 µm coronal sections and stored in cryoprotectant at -20 °C or 10 µm sections and directly mounted on positively charged slides.

#### Fresh-frozen

Hippocampus and corpus callosum tissue was dissected from 3-, 9-, and 15-month wildtype and APP/PS1 rats, distinct from the perfused animals described above. From each brain region, one tissue punch was flash frozen on dry ice for protein isolation, and another was placed in 500 µL of TRIzol (Life Technologies) for RNA isolation. All samples were stored at -80 °C until further processing.

### 2.5 Nanoscale Flow Cytometry

Isolated plasma was thawed once at room temperature and separated into 50 µl aliquots before refreezing at -80 °C, samples were all run on their second thaw cycle. Plasma aliquots were thawed at room temperature prior to incubating 10 µl with anti-TMEM119 Coralite-647 (200 ng/sample, Proteintech 66948) and anti-TREM2 Alexa Fluor-405 (400 ng/sample, Novus Biologicals 07101) for 30 mins at room temperature in the dark. After incubation, samples were diluted 100-fold with PBS (final dilution of 200-fold) in a final concentration of 0.0125% Triton-X-100 to permeabilize EVs. Triplicates of each incubation were run on the Apogee A50 Microplus Nanoflow Cytometer (Apogee Flow Systems Inc). Background levels of PBS were below 200 events/s and standardized beads were measured at their stock concentration of 5000 events/s prior to running plasma samples. The following instrument settings were kept consistent across all samples: flow rate: 150 µL/min for 120 µL, sheath pressure: 150 mbar, lasers: 50 mW 405 nm, 50 mW 638 nm, 50 mW 488 nm, photomultiplier (PMT) voltages: SALS (350 V), LALS (300 V), L405-Blu (450 V), and L638-Red (475 V). Thresholds to eliminate background noise were set at 3 a.u. for small angle light scatter (SALS) and 20 a.u. for long angle light scatter (LALS). Fig. S2 displays polystyrene and silicon beads for comparison of EV sizes, as well as unlabelled plasma, dilution reagent (PBS), and individual antibodies. Nanoscale flow cytometry data is reported as events/µL and indicates the concentration of labelled particles detected after gating for TMEM119 and TREM2 fluorescent antibodies.

### 2.6 EV isolation and immunoprecipitation

#### EV isolation

Plasma was thawed at room temperature and EVs were isolated from 1 mL of rat plasma via 35 nm pore size-exclusion chromatography columns (IZON Science). EVs were then concentrated with centrifugation filters (Amicon Ultra-15) prior to western blotting.

#### Immunoprecipitation

Protein concentration of the SEC-isolated EVs was measured using a BCA assay prior to incubation with biotinylated TMEM119 (Novus Biologicals 10313B) (100 µg of EVs with 1:50 TMEM119 diluted in PBS) overnight at 4 °C with rotation. The following day samples were incubated with 40 µL of streptavidin magnetic beads (Thermo Fisher Scientific 65601) for 30 mins at room temperature with rotation. The tubes were then placed on a magnetic rack (Thermo Fisher Scientific), liquid was pipetted off, and the beads were washed 4 times with 200 µL of PBS prior to sample lysis in 50 µL of 2% SDS heated at 95 °C for 15 minutes. After cooling at room temperature for 5 minutes, tubes were placed back on the magnetic rack and supernatant was retained.

### 2.7 RNA isolation and qPCR

#### RNA isolation

Tissue samples from the hippocampus and corpus callosum were brought up to 600 µL of TRIzol and mechanically homogenized. Samples were vortexed with chloroform (120 µL) and centrifuged at 12,000 x *g* for 15 mins at 4 °C. The aqueous phase was pipetted off and mixed with an equal volume of isopropyl alcohol prior to incubation at -20 °C for 30 mins. Following incubation, samples were centrifuged at 12,000 x *g* for 10 mins at 4°C. The supernatant was then removed, and the remaining pellet was washed with 75% ethanol twice by centrifugation at 7,500 x *g* for 5 mins at 4 °C. The pellet was air-dried and resuspended in 15 µL of RNase-free water prior to determining RNA concentrations using a Nanodrop One spectrophotometer (Thermo Fisher Scientific).

#### qPCR

Specific forward and reverse primers were designed using the NCBI primer design tool and provided in Table S1. Primers were combined with 2 µL of cDNA and SsoAdvanced Universal SYBR Green Mix (Bio-Rad). RPL13⍰ and β-actin housekeeping genes were averaged and together used as an endogenous control. All mRNA expression levels were normalized to this value and comparison of transcript levels were performed using the Δ ΔC^T^ method^42^.

### 2.8 Protein isolation and western blot

#### Protein isolation

Lysis buffer (150 mM NaCl, 1.0% IGEPAL CA-630, 0.5% sodium deoxycholate, 0.1% SDS, 50 mM Tris, pH 8.0) was added to tissue samples collected from the hippocampus and corpus callosum and tissue was mechanically homogenized. Tissue lysates were sonicated and then centrifuged at 15,000 x *g* for 15 mins at 4 °C. The supernatant was retained, and protein concentration was determined using a BCA protein assay kit (Pierce, Thermo Fisher Scientific) according to the manufacturer’s instructions.

#### Western blot

Protein was diluted in loading buffer (1 x LDS, 5mM DTT in 0.5% SDS) and denatured for 10 mins at 70 °C. Protein separation was performed using gel electrophoresis Bis-Tris acrylamide gels in MOPS SDS running buffer (Thermo Fisher Scientific) at 70 mA per gel. Transfer to PVDF membrane (Roche Diagnostics) was performed at 100 V for 100 minutes on ice and then membranes were blocked with 5% BSA in Tris Buffered Saline-Tween (TBST) (50 mM Tris-HCl, pH 8.0, 0.15 M NaCl, 0.1% Tween) overnight at 4 °C. The next day, the membranes were incubated with primary antibodies in 5% BSA in TBST for 1 hour at room temperature (TMEM119; 1:1000, Proteintech 66948 and ⍰-tubulin; 1:1000, Santa Cruz 5286) or overnight at 4 °C (TREM2; 1:1000, Bioss 2723R). Membranes were washed for 3 x 10 mins with TBST prior to incubation with anti-mouse (TMEM119, ⍰-tubulin) or anti-rabbit (TREM2) HRP-conjugated secondary antibodies (1:10,000, Jackson Laboratories) for 1 hour at room temperature. Membranes were washed again for 3 x 10 mins prior to visualization with chemiluminescent HRP substrate (Immobilon) and imaging using a ChemiDoc MP system (Bio-Rad). The volume tool function in ImageLab (BioRad) was used to quantify each band of interest and protein expression was normalized to ⍰-tubulin expression.

### 2.9 Histology

#### Immunohistochemistry

A diaminobenzidine (DAB) staining protocol was followed on free-floating 30 µm sections from bregma levels +2.00 mm and -3.00 mm. Tissue sections were first rinsed in 0.01 M PBS for 6 x 10 mins and then incubated with 1% H_2_0_2_ (Thermo Fisher Scientific) for 15 mins. Next, sections were incubated with primary antibody against TREM2 (1:1000, R&D Systems AF1729) in blocking solution (2% rabbit serum in PBS with 0.4% Triton-X 100) for 1 hour at room temperature and 48 hours at 4 °C. The next day, sections were incubated with biotinylated rabbit anti-sheep secondary antibody (1:500, Vector Laboratories VECTBA6000) in blocking solution for 1 hour at room temperature, processed using an ABC kit (Thermo Fisher Scientific), and visualized using DAB (Sigma-Aldrich), with 0.01 M PBS washes between each step. Stained sections were mounted onto slides using 0.3% gelatin and airdried overnight. Lastly, slides were dehydrated using progressive concentrations of ethanol and xylene and then cover slipped with Depex mounting medium (Electron Microscopy Sciences). For each round of staining, sections from all experimental groups were processed in parallel using the same solutions.

#### Immunofluorescence

Pre-mounted 10 µm sections were first washed for 3 x 5 mins with 0.01 M PBS and then incubated overnight at 4 °C with primary antibodies against TMEM119 (1:500, Proteintech CL-64766948) and TREM2 (1:200, R&D Systems AF1729) in 3% BSA and 0.3% Triton-X-100 in PBS. The following day the sections were washed with PBS for 3 x 5 mins and then incubated with donkey anti-sheep 488 conjugated secondary antibody (1:500, Thermo Fisher A-11015) for 1 hour at room temperature. Lastly, slides were washed with PBS for 3 x 5 mins prior to cover slipping with DAPI mounting medium.

### 2.10 Microscopy & image analysis

All image analysis was conducted by an experimenter blinded to the treatment groups. Images were taken using an upright brightfield microscope (Nikon Eclipse Ni-E, NIS Elements) or an inverted microscope (Mica Microhub, Leica Microsystems). Example images were captured using 10 ⍰ and 20 ⍰ objectives. TREM2 quantification was carried out on six images randomly captured at 40 ⍰ in the hippocampus (bregma -3.00 mm) and corpus callosum (bregma +2.00 mm). White balance was automated, and settings were kept consistent for all images. Cell density was assessed using the Fiji ImageJ software^43^ cell counter plugin in which TREM2^+^ cells were manually selected in each image. The number of TREM2^+^ cells was averaged across the six images and divided by the area to obtain the TREM2^+^ cell density in each region of interest.

### 2.11 Data analysis

Statistical analyses were conducted using SPSS (IBM Corporation) and GraphPad Prism software. Depending on the comparison, unpaired *t*-tests, two-way, or three-way analysis of variances (ANOVA) were performed with a significance value of *p* = 0.05 and when required, Bonferroni’s *post-hoc* correction was used. When Mauchly’s test of sphericity was violated in the repeated-measures ANOVA, the Greenhouse-Geisser correction was applied. Welch’s correction was used on *t*-tests with unequal variances between groups. Methodology schematics were generated using BioRender (Biorender.com) and graphs were created using Graphpad Prism. All data are presented as mean values with error bars indicating standard error of the mean (SEM).

## 3. Results

### 3.1 Increased circulating TMEM119^+^/TREM2^+^ EVs in 15-month-old APP/PS1 rats

Previous work has identified TREM2 as a microglial marker critical in AD progression, thus, we sought to identify TREM2^+^ MEVs in the systemic circulation as a potential biomarker of brain TREM2 expression. To achieve cell specificity, we dual-labelled with microglial transmembrane protein 119 (TMEM119), previously shown to differentiate microglia from macrophages^44–47^.

Using nanoscale flow cytometry, we analyzed TMEM119^+^/TREM2^+^ EVs in 3-, 9-, and 15-month-old WT and APP/PS1 rat plasma. Fig. 1A shows example scatter plots of TMEM119, TREM2, and dual-positive particles labelled directly in rat plasma. TMEM119^+^/TREM2^+^ events/µL were significantly increased in 15-month-old APP/PS1 rats compared to wildtypes (Fig. 1B; main effect of genotype: F_(1, 41)_ = 9.261, *p* = 0.004, η_p_^2^= 0.184). There was no significant difference in dual-labelled events/µL between genotypes at 3- or 9-months of age and no sex differences were observed at any age, thus males and females were collapsed (Fig. 1B).

**Figure 1:**
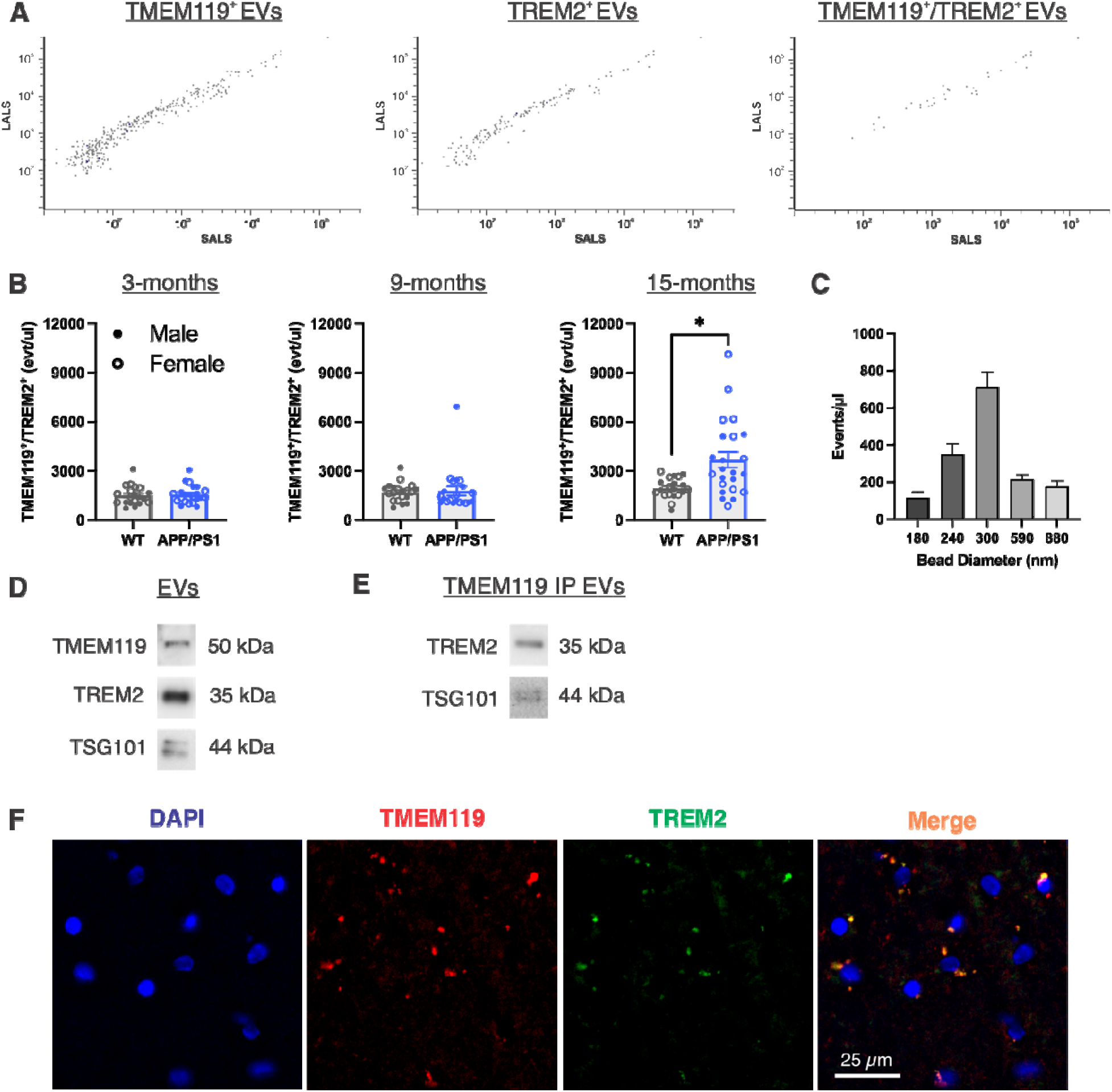
TMEM119^+^/TREM2^+^ EVs are increased in 15-month-old APP/PS1 rats. A) Example nanoscale flow cytometry scatter plots of TMEM119, TREM2 and dual-positive EVs. B) TMEM119^+^/TREM2^+^ events/µl labelled in 3-, 9-, and 15-month wildtype and APP/PS1 rat plasma assessed using a 2-way ANOVA (genotype ⍰ sex). C) Size distribution of TMEM119^+/^TREM2^+^ EVs based on standardized reference beads. D) Western blot of TMEM119 and TREM2 from isolated rat plasma EVs. E) Western blot of TREM2 from immunoprecipitated TMEM119^+^ EVs. F) Immunofluorescent 20 ⍰ images of TMEM119 and TREM2 in 15-month-old APP/PS1 hippocampus. Scale bar indicates 25 µm. * Indicates statistical significance (*p* < 0.05) Data represent the group mean ± SEM. *n* = 10-12 males and females per group. LALS, long angle light scatter; SALS, short angle light scatter.

To further evaluate the ability to separate APP/PS1 rats from wildtypes, receiver operating characteristic (ROC) curves were generated for TMEM119^+^/TREM2^+^ EVs at each age. The analysis revealed limited discrimination between genotypes at 3- and 9-months, but the 15-month ROC showed moderate discriminability with an area under the curve (AUC) of 0.757 (sensitivity, 69.57%; specificity, 75.00%) (Fig. S3). Based on standardized reference beads, a size distribution of dual-labelled TMEM119^+^/TREM2^+^ EVs showed the largest proportion of EVs falling in the 300 nm range (Fig. 1C). Using SEC and immunoblotting we showed the presence of TMEM119 and TREM2 proteins in plasma EVs (Fig. 1D). Following immunoprecipitation with an antibody against TMEM119, we revealed enrichment of TREM2 protein in isolated TMEM119^+^ EVs (Fig. 1E). Immunofluorescence confirmed microglia co-expressed TMEM119 and TREM2 in APP/PS1 rat brain tissue (Fig. 1F). Overall, TREM2^+^ MEVs were effectively detected and validated in the systemic circulation, and were elevated in 15-month APP/PS1 rats.

### 3.2 Anatomical, age, and genotype-dependent differences in brain TREM2 expression

Hippocampal levels of TMEM119 and TREM2 expression were measured using qPCR and western blot. The hippocampus was chosen as a region of interest (Fig. 2A) due to the known association of TREM2^+^ microglia with Aβ^48,49^, and location of plaque deposition in this model (Fig. S1). TMEM119 gene expression was increased between 3- and 9-month rats but no genotype-related differences were observed (Fig. 2B; main effect of age: F_(2, 25)_ = 3.617, *p* = 0.042, η_p_^2^ = 0.224, *p*_bonf_ = 0.042). Likewise, TREM2 gene expression was unchanged between genotypes but was increased at 15-months compared to 3- and 9-month rats (Fig. 2B; main effect of age: F_(2, 26)_ = 9.629, *p* = <0.001, η_p_^2^ = 0.426, p_bonf_ = 0.009). Western blot detection of TMEM119 protein levels showed no difference in TMEM119 abundance between genotypes at any age timepoint (Fig. 2C-E). Further, wildtype and APP/PS1 rats had similar TREM2 protein levels at 3-months (Fig. 2C), but in contrast to our qPCR results, TREM2 protein was significantly increased in APP/PS1 rats at 9-months (Fig. 2D; t_(10)_ = 5.533, *p* = 0.0003) and significantly decreased in APP/PS1 rats at 15-months compared to the wildtypes (Fig. 2E; t_(10)_ = 4.791, *p* = 0.0007). Given the demonstrated increase in white matter microglia activation in aging and AD^5,50^, and the identification of TREM2 as a WAM marker^15^, we next analyzed TMEM119 and TREM2 expression in the corpus callosum (Fig. 3A). No differences in TMEM119 gene expression were observed but TREM2 was upregulated at 15-months compared to 3- and 9-month rats (Fig. 3B; main effect of age: F_(2, 27)_ = 20.63, *p* = <0.001, η_p_^2^ = 0.604, *p*_bonf_ = <0.001). Western blot analysis revealed no genotype-related changes in TMEM119 or TREM2 protein expression at any age in the corpus callosum (Fig. 3C-E).

**Figure 2:**
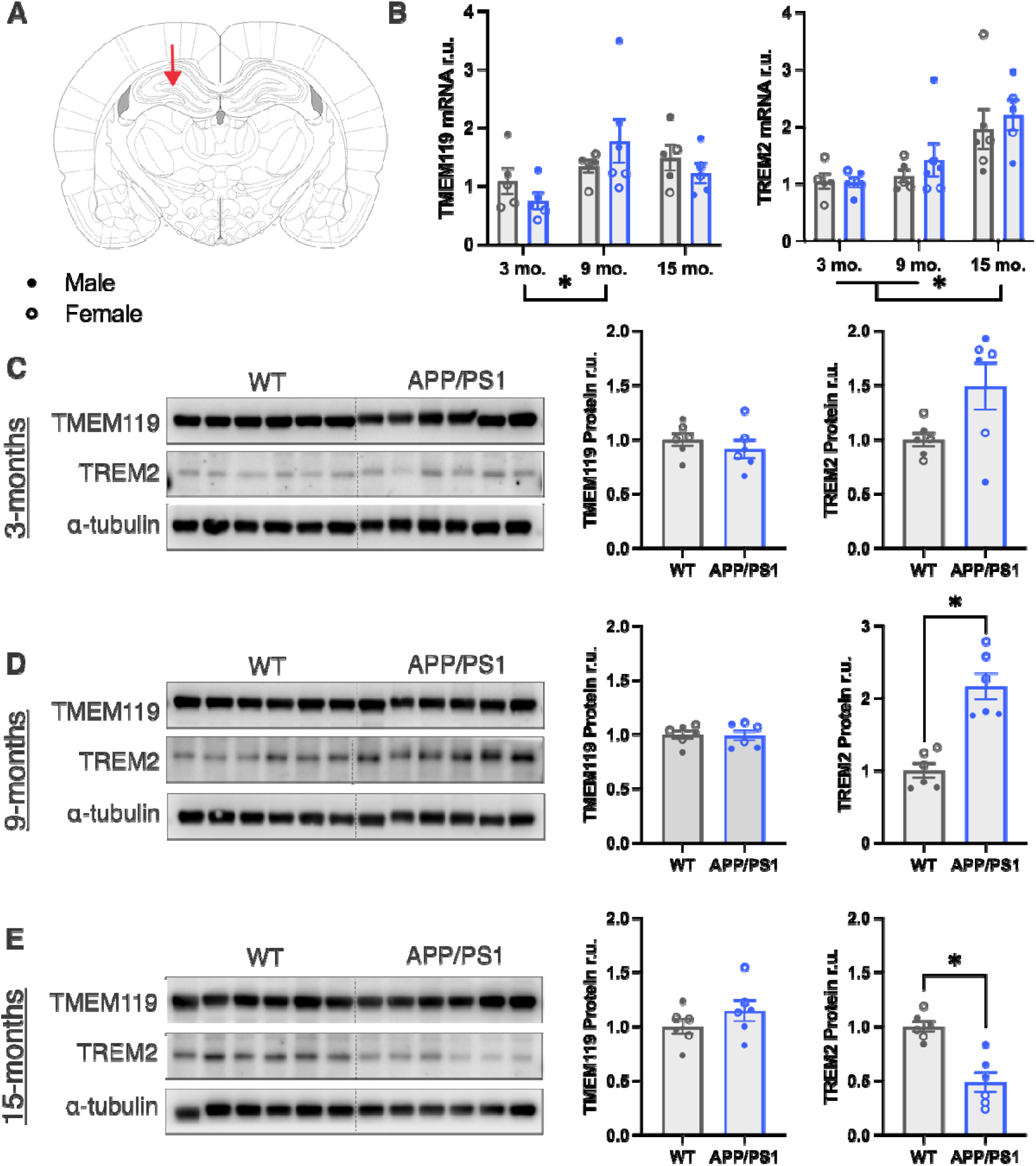
TREM2 mRNA and protein levels differ by age and genotype in the hippocampus. A) Coronal section for the hippocampal region of interest. B) qPCR measurement of TMEM119 and TREM2 in 3-, 9-, and 15-month WT and APP/PS1 hippocampal RNA isolates assessed using 2-way ANOVAs (age ⍰ genotype). C-E) Western blot of TMEM119 and TREM2 and corresponding quantification in hippocampal protein isolates from 3-, 9-, and 15-month WT and APP/PS1 rats assessed using *t*-tests. * Indicates statistical significance (*p* < 0.05). Data represent the group mean ± SEM. *n* = 2-3 males and 2-3 females per group.

**Figure 3:**
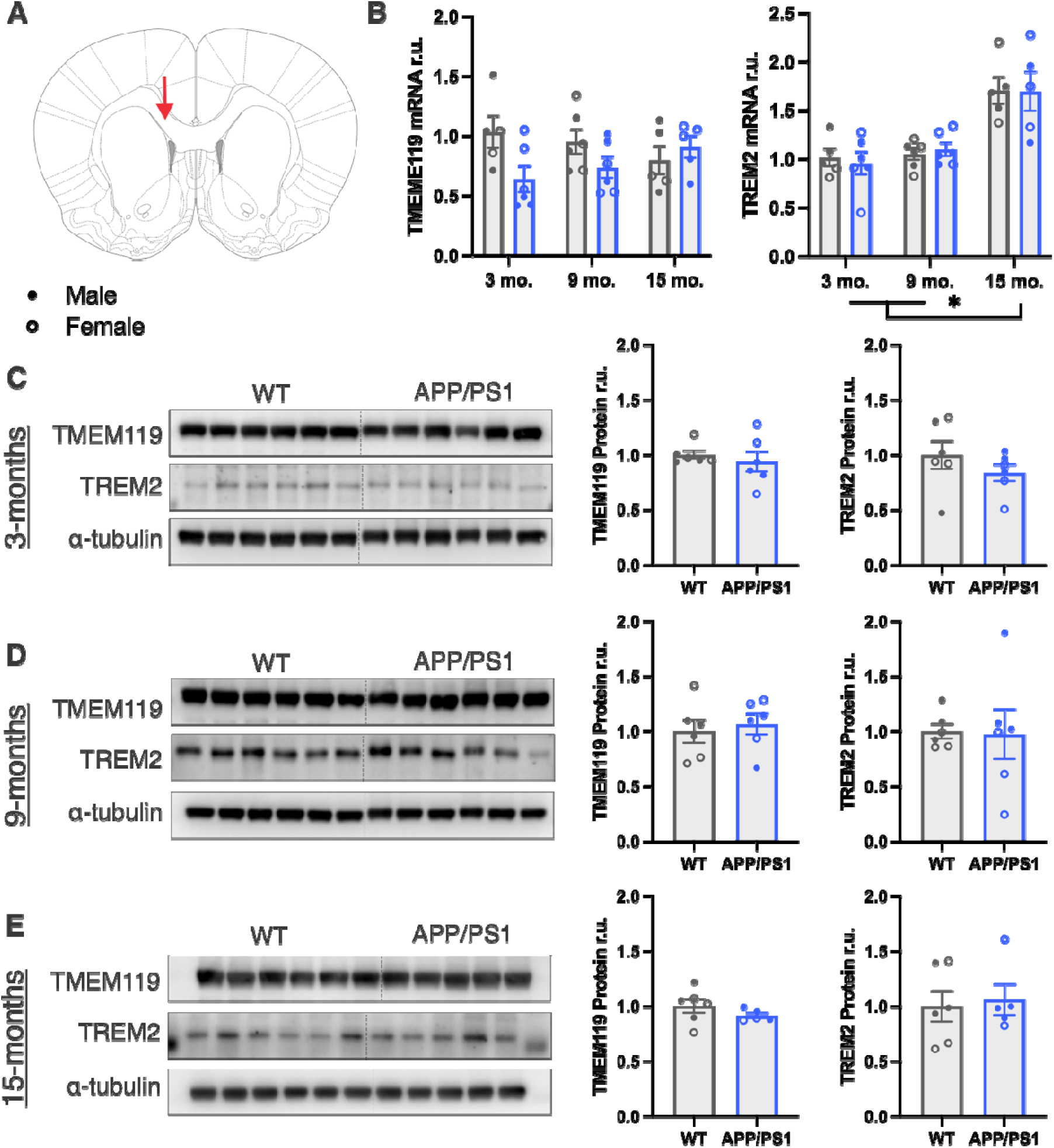
Increased TREM2 mRNA in the corpus callosum of 15-month-old rats. A) Coronal section for the corpus callosum region of interest. B) qPCR measurement of TMEM119 and TREM2 in 3-, 9-, and 15-month WT and APP/PS1 corpus callosum RNA isolates assessed using 2-way ANOVAs (age ⍰ genotype). C-E) Western blot of TMEM119 and TREM2 and corresponding quantification in corpus callosum protein isolates from 3-, 9-, and 15-month WT and APP/PS1 rats assessed using *t*-tests. * Indicates statistical significance (*p* < 0.05). Data represent the group mean ± SEM. *n* = 2-3 males and 2-3 females per group.

Given the contrasting results in TREM2 gene and protein expression, we next conducted IHC to label for TREM2^+^ cells in the hippocampus (Bregma -3.00 mm) and corpus callosum (Bregma +2.00 mm). In the hippocampus, we found increased TREM2^+^ cell density in 9- and 15-month-old APP/PS1 rats compared to wildtypes (Fig. 4A; 9-months: t_(14)_ = 2.901, *p* = 0.0116; 15-months: t_(14)_ = 4.972, *p* = 0.0002). In the corpus callosum, TREM2 cell density was unchanged at 3-, and 9-months, but was increased in 15-month APP/PS1 rats compared to wildtypes (Fig. 4C; t_(9.677)_ = 3.76, *p* = 0.0039). Altogether, we revealed complexity in TREM2 brain expression in this model which differed by neuroanatomical area (i.e., hippocampus vs. corpus callosum) and method of detection (i.e., qPCR vs. western blotting vs. IHC).

**Figure 4:**
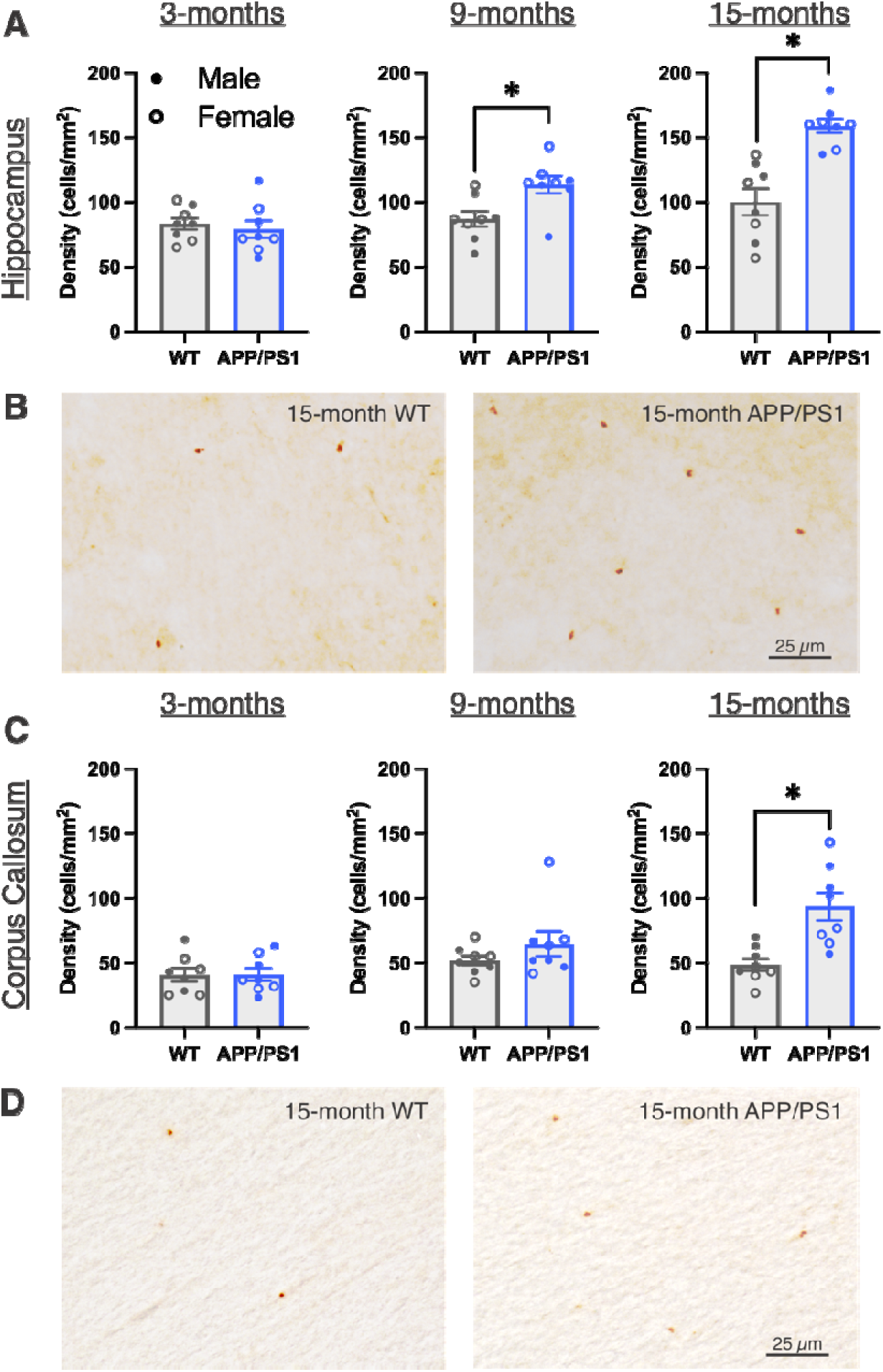
Increased histological TREM2 expression in aged APP/PS1 rats. A) Quantification of TREM2 cell density in the hippocampus of 3-, 9-, and 15-month WT and APP/PS1 rats assessed using *t*-tests. B) Representative 20 ⍰ images of TREM2^+^ cells in the hippocampus of 15 month WT and APP/PS1 rats. C) Quantification of TREM2 cell density in the corpus callosum of 3-, 9-, and 15-month WT and APP/PS1 rats assessed using *t*-tests. D) Representative 20 ⍰ images of TREM2^+^ cells in the corpus callosum of 15-month WT and APP/PS1 rats. Scale bar indicates 25 µm. * Indicates statistical significance (*p* < 0.05). Data represent the group mean ± SEM. *n* = 4 males and 4 females per group.

### 3.3 Impairment in spatial working and reference memory associates with amount of circulating TMEM119^+^/TREM2^+^ EVs in 15-months-old rats

To determine the effects of age, genotype, and sex on cognition in our model, rats were tested in a 4/8 RAWM to assess spatial working and reference memory (Fig. 5A). Errors were summed in three-day increments at each age for both spatial reference and working memory (Fig. 5B and C). Rats at all ages demonstrated improved spatial memory, as shown by decreased errors across the 15 testing days in both reference memory (main effect of day for all ages; 3-months: F_(4, 148)_ = 107.445, *p* < 0.001, η_p_^2^ = 0.744; 9-months: F_(3.166, 113.96)_ = 48.307, *p* < 0.001, η_p_^2^ = 0.573; 15-months: F_(3.239, 126.311)_ = 30.02, *p* < 0.001, η_p_^2^ = 0.435) and working memory (main effect of day for all ages; 3-months: F_(3.102, 114.786)_ = 159.698, *p* < 0.001, η_p_^2^ = 0.812; 9-months: F_(3.065, 110.34)_ = 49.736, *p* < 0.001, η_p_^2^ = 0.58; 15-months: F_(4,156)_ = 29.198, *p* < 0.001, η_p_^2^ = 0.428). Spatial reference memory was impaired in 3-month APP/PS1 rats as they committed more errors than wildtypes on days 6, 9, and 12, but reached the same level of proficiency by day 15 (Fig. 5B; genotype ⍰ sex ⍰ day interaction; F(4, 148) = 3.59, *p* = 0.008, η_p_^2^ = 0.088; day 6 WT vs APP/PS1, *p*_bonf_ = 0.003; day 9 WT male vs APP/PS1 male, *p*_bonf_ < 0.001; day 12 WT vs APP/PS1, *p*_bonf_ = 0.004). Moreover, APP/PS1 rats committed more reference memory errors overall compared to wildtypes at both 9- and 15-months (Fig. 5B; main effect of genotype; 9-months: F_(1, 36)_ = 4.458, *p* = 0.042, η_p_^2^ = 0.11; 15-months: F_(1, 39)_ = 30.933, *p* < 0.001, η_p_^2^ = 0.442). The only sex-specific result observed in this task occurred in the 3-month reference memory errors on day 9 in which the difference was specifically between the wildtype male and APP/PS1 male rats (day 9 WT male vs APP/PS1 male, *p*_bonf_ < 0.0001). Spatial working memory impairment was observed later in this model in which wildtype and APP/PS1 rats performed similarly at 3- and 9-months-old but APP/PS1 rats committed more spatial working memory errors overall at 15-months (Fig. 5C; main effect of genotype; F_(1, 39)_ = 25.788, *p* < 0.001, η_p_^2^ = 0.398). To assess cumulative performance on the task, we totalled all errors committed across the 15 testing days. APP/PS1 rats committed significantly more total errors than wildtypes at 3- and 15-months, but not 9-months, with no sex-specific differences (Fig 5D; main effect of genotype; 3-months: F_(1, 37)_ = 8.283, *p* = 0.007, η_p_^2^ = 0.183; 15-months: F_(1, 39)_ = 37.619, *p* < 0.001, η_p_^2^ = 0.491). To investigate the potential relationship between extent of cognitive impairment and circulating MEVs in aged rats, we performed linear regressions at each age between total errors and TMEM119^+^/TREM2^+^ events/µl and ultimately found a significant association in 15-month-old rats (Fig. 6A; *p* = 0.031, r = 0.33, R^2^ = 0.11). Taken together, our novel investigation not only revealed a progressive, age-related and APP/PS1-linked impairment of spatial working and reference memory, but this cognitive impairment of the 15-month-old rats was also associated with their levels of circulating TMEM119^+^/TREM2^+^ EVs; important findings when considering MEVs as a potential biomarker for AD-related cognitive decline across aging.

**Figure 5:**
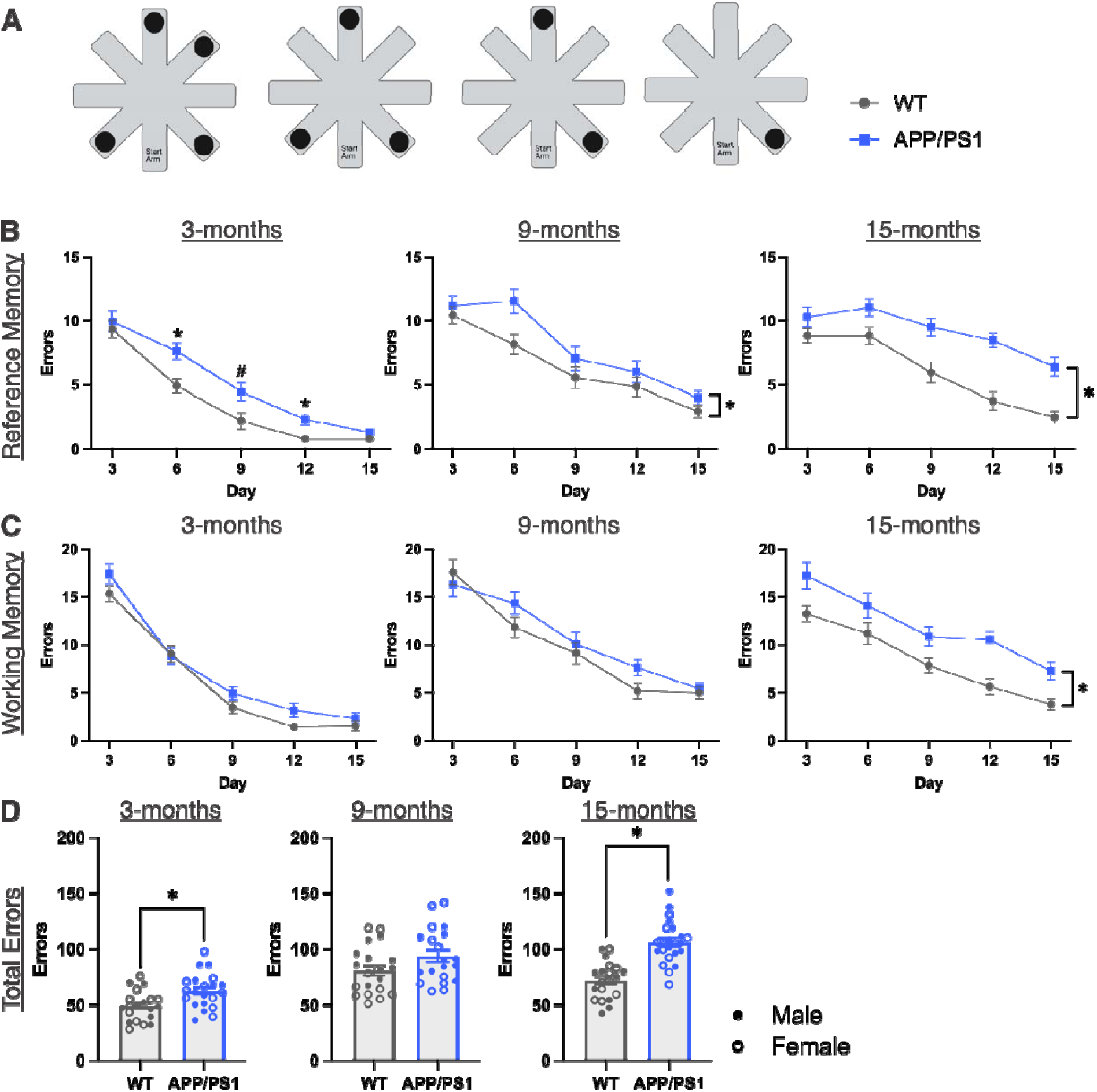
Spatial working and reference memory impairments in APP/PS1 rats. A) Schematic of four daily RAWM trials in an 8-arm maze with 4 platforms. B) Reference memory errors committed across the 15 testing days displayed as sums per 3-day increment in 3-, 9-, and 15-month WT and APP/PS1 rats assessed using 3-way mixed ANOVAs (genotype ⍰ sex ⍰ day). C) Working memory errors committed across the 15 testing days displayed as sums per 3-day increment in 3-, 9-, and 15-month WT and APP/PS1 rats assessed using 3-way mixed ANOVAs (genotype ⍰ sex ⍰ day). D) Total errors committed in the RAWM task in 3-, 9-, and 15-month WT and APP/PS1 rats assessed using *t*-tests. * Indicates statistical significance (*p* < 0.05) between WT and APP/PS1 groups. # Indicates statistical significance (*p*_bonf_ < 0.05) between WT males and APP/PS1 males. Data represent the group mean ± SEM. *n* = 10-12 males and females per group. Schematic created with Biorender.com

**Figure 6.**
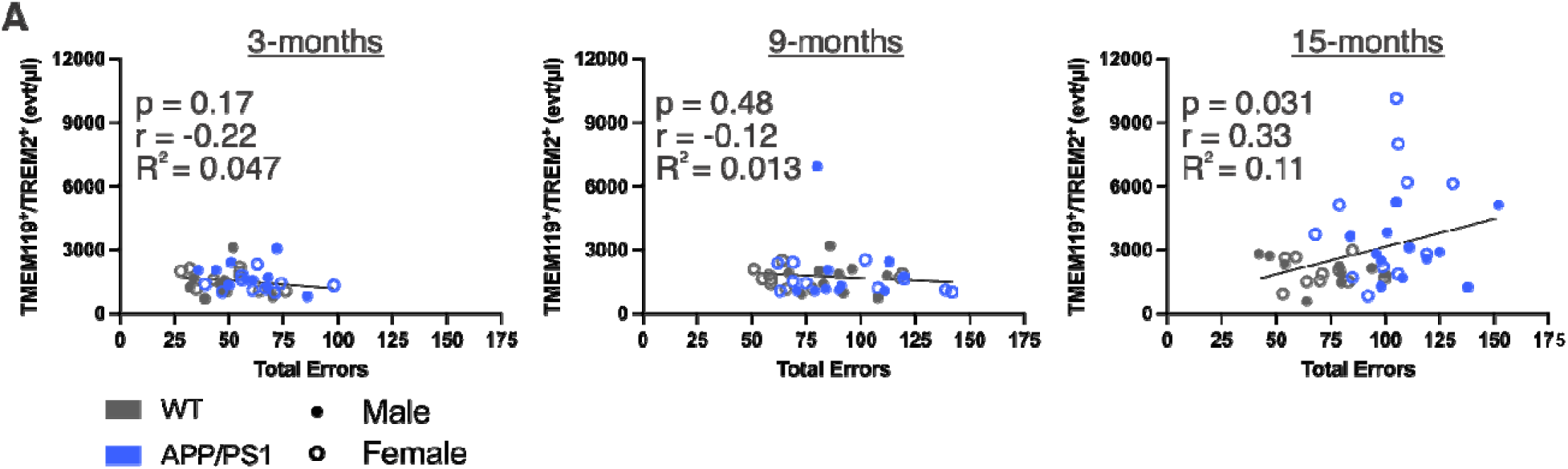
Impaired spatial memory associates with increased TMEM119^+^/TREM2^+^ EVs in 15-month-old rats. A) Simple linear regression between TMEM119^+^/TREM2^+^ events/µl and total RAWM errors in 3-, 9-, and 15-month WT and APP/PS1 rats. *n* = 10-12 males and females per group.

## 4. Discussion

Microglia play critical roles in AD pathogenesis, but we lack non-invasive and sensitive methods to measure them *in vivo*. In the present study, the primary objective was to investigate plasma-derived MEVs as a biomarker of brain microglial expression in a rat model of aging and AD. We hypothesized that given the previously described increase in DAMs and WAMs (TREM2^+^) in AD^15,17^, we would observe an increase in circulating EVs with this signature. In line with our prediction, TMEM119^+^/TREM2^+^ EVs were increased in aged APP/PS1 rats. Interestingly, TREM2 expression in the brain was complex, such that the results depended on age, method of detection, and anatomical region. Additionally, we sought to evaluate cognition and found that level of impairment associated with amount of TMEM119^+^/TREM2^+^ EVs. To our knowledge, this is the first study to detect TREM2^+^ MEVs in plasma and investigate the relationship between circulating MEVs and cognition in AD.

### 4.1 Circulating TMEM119^+^/TREM2^+^ EVs as a biomarker of brain microglial phenotype

Since circulating EVs originate from cells throughout the body, and TREM2 is not only expressed by microglia, we dual-labelled EVs with microglial marker TMEM119, and assessed its expression in the same brain regions. Previous studies have validated TMEM119 as a marker that differentiates microglia from macrophages and is expressed on microglial EVs^44–47^.

Downregulation of TMEM119 gene and protein expression has been described in reactive microglia and DAMs^17,51,52^, challenging its use as a robust marker of microglia under pathological conditions. In human AD brain tissue, a decrease in microglia expressing TMEM119 has been observed compared to age-matched controls^53,54^ and in small EVs isolated from human AD tissue^55^. Importantly, and in contrast to these studies, our results showed no genotype-related differences in TMEM119 expression in the corpus callosum or hippocampus. The consistency in TMEM119 expression between wildtype and APP/PS1 brain tissue suggests that it served as a suitable microglial marker in this model, but further work is needed to consider species-related and region-specific differences in TMEM119 expression in AD.

Given that recent studies indicate TREM2 is important in AD risk and pathogenesis, but *in vivo* measurement is challenged by a lack of specificity, we set out to detect circulating TREM2^+^ MEVs. Limited findings on TREM2^+^ EVs have been described in isolations from AD human brain tissue and *in vitro* models^55–57^, including an increase in TREM2^+^ EVs isolated from human AD parietal cortex^55^. In line with this, we observed an increase in circulating TMEM119^+^/TREM2^+^ EVs in 15-month-old APP/PS1 rats. As ROC analysis of the 15-month TMEM119^+^/TREM2^+^ EVs also showed moderate discrimination between genotypes, future studies should explore combining this with other AD blood-based biomarkers (i.e., Aβ, tau) for improved discrimination. The simultaneous loss of brain TREM2 protein and increase in circulating TREM2^+^ MEVs at 15-months might be explained by increased shedding of TREM2 protein via EVs. Although further experiments are needed to explore this in TREM2^+^ microglia, increased EV shedding and altered proteomic composition have been previously described under cellular stress conditions^58,59^. Rate of EV release may also be influenced by factors such as ATP and pro-inflammatory molecules^60,61^, an important consideration in better understanding circulating MEVs as they relate to brain microglial activity in AD.

EVs encompass multiple subpopulations, including exosomes, microvesicles, and apoptotic bodies, which differ in size and release pathways. While we can’t confirm subpopulation of EVs in this study, the size distribution (majority in the 300 nm range) determined by light-scatter based measurements of size-standardized beads suggests the TMEM119^+^/TREM2^+^ EVs detected were larger than exosomes. Our size distribution falls within a similar range as CD14^+^ MEVs and large MEVs previously described in an experimental model of stroke and an *in vitro* AD model^44,62^. For specific measurement of TMEM119^+^/TREM2^+^ EV size, isolation via immunoprecipitation followed by fluorescent nanoparticle tracking can be used in the future. Further protein analyses could also help confirm the main EV subpopulation and the corresponding release pathway. This will be critical in understanding how EV subtype could affect the temporal profile of EV release in the brain vs detection in the systemic circulation, timing that is not yet understood.

### 4.2 TREM2 brain expression in the APP/PS1 rat model

Previous investigations into TREM2 and its role in AD have produced varying results, thus, it was important to characterize TREM2 expression in this model across age, using multiple techniques. While studies have shown increased TREM2 protein and RNA in mouse and clinical AD brain samples ^13,17,63^, some studies have also observed downregulation in TREM2 at the protein level, possibly dependent on stage of disease and brain region^64–66^. Consistent with the idea that TREM2 expression depends on disease stage, we found that hippocampal TREM2 protein levels were increased in 9-month-old APP/PS1 rats but decreased in 15-month-old APP/PS1 rats, despite no genotype-related change in TREM2 RNA levels. This was further complicated by an increase in TREM2^+^ cells in the hippocampus of 9- and 15-month-old APP/PS1 rats detected via immunohistochemistry. Since Aβ plaque load is low in the 9-month rats (Fig. S1), perhaps this increase at 9 months represents functional TREM2^+^ phagocytosing microglia. In contrast, the increase in hippocampal TREM2^+^ cells but decrease in TREM2 protein at 15 months possibly indicates a reduction in functional TREM2 protein levels as Aβ plaques accumulate. This is supported by evidence that microglial genes, including TREM2, undergo alternative splicing in AD, leading to reduced protein expression and altered functions^67,68^. Overall, our results support the idea that there is variation in TREM2 brain expression contingent on age, anatomical region, species, and mode of detection and caution against relying exclusively on RNA sequencing for microglial phenotyping.

### 4.3 TMEM119^+^/TREM2^+^ EVs associate with the severity of cognitive impairment

Our next objective was to characterize cognition in the APP/PS1 rats across age. Previous testing in this model found spatial learning and memory deficits in the APP/PS1 rats using the Barnes maze in 7-, 10-, 12-, and 14-month-old rats across multiple studies^39,69,70^. We employed a RAWM, which assessed both spatial reference and working memory. The previous finding of spatial reference memory deficits as early as 7-months-old^69^, aligns with the reference memory impairments we observed in the 9- and 15-month APP/PS1 rats. We also showed minor reference memory deficits prior to onset of Aβ plaque deposition in the 3-month APP/PS1 rats, highlighting the need for further assessment of early pathologies. Based on experimental and clinical studies, executive function deficits have been observed in mild cognitive impairment and early in AD progression^5,71,72^. However, we found that reference memory impairment preceded working memory impairment (a subdomain of executive function) in the APP/PS1 rats. Investigating additional executive function subdomains (i.e., cognitive flexibility; response inhibition) could reveal intricacies in subdomain trajectories in aging and AD. Ultimately, the RAWM allowed for assessment of cognition that aligned with previous testing in APP/PS1 rats and added to our understanding of multiple cognitive domains in this model.

Lastly, we aimed to assess the relationship between cognition and TMEM119^+^/TREM2^+^ EVs. To do this, we summed the total errors committed in the RAWM task, thereby providing a more cumulative assessment of rodent cognition across multiple domains, and ultimately found a significant positive association between total errors and circulating TMEM119^+^/TREM2^+^ EVs in 15-month-old rats. Previous investigation into TREM2 as a biomarker has focused on its soluble form (sTREM2), with evidence that sTREM2 in cerebrospinal fluid rises in early AD but declines later in disease progression^73,74^. However, to our knowledge, our results represent the first plasma-based, cell-specific evidence of TREM2 as a biomarker of cognition. The relationship between cognitive performance and TMEM119^+^/TREM2^+^ EVs exhibited a relatively low R^2^ value (0.11), suggesting limited predictive power; however, this could be related to the observed complexity of TREM2 expression levels, as discussed above. Looking forward, given that the APP/PS1 rats demonstrated impaired cognition as early as 3 months, future assessment of MEVs as biomarkers could explore a combination of microglial markers and cargo that may change earlier in AD progression.

Given the increased risk for AD in females, it was important to consider sex-related differences. In line with a previous report of no cognitive differences between male and female 10-month APP/PS1 rats in the Barnes maze^39^, we did not observe sex differences in the RAWM task, aside from a singular day in the 3-month rats in which the APP/PS1 males differed from the wildtype males but not females. There is also evidence of sexual dimorphism in microglial expression and EV signatures^75,76^, and while there were balanced numbers of males and females, we were not powered to include sex in the statistical analyses for IHC, protein, or RNA. However, the EV analysis revealed no sex-related differences in the amount of circulating TMEM119^+^/TREM2^+^ EVs. Future inclusion of sex as a factor will give more insight into sex differences in microglial phenotypes.

In conclusion, we detected TMEM119^+^/TREM2^+^ EVs in plasma for the first time using nanoscale flow cytometry, providing insight into their potential use as a biomarker of brain TREM2 expression and cognitive impairment. Further experiments are needed to better understand the relationship between brain microglia and circulating MEVs, cause of increased TREM2^+^ EV release, and how this affects cellular function. TREM2 poses an interesting therapeutic target in AD and other neurological conditions, but more work is required to understand its role in different stages of disease progression. The measurement of TMEM119^+^/TREM2^+^ EVs using nanoscale flow cytometry offers a non-invasive approach to longitudinal tracking of TREM2, with possible application to various age-related neurological conditions.

## Acknowledgements

We would like to thank Dr. Lynn Wang, Dr. Lin Zhao, and UWO’s Animal Care and Veterinary Services staff for their technical assistance.

## Author contribution

SJM conducted rodent cognitive testing. SJM and ADR performed plasma/tissue collections, molecular experiments, and data analysis. SJM completed manuscript writing. All authors contributed to research design, interpretation and manuscript editing.

## Data Availability

Data will be made available on request.

---

**Table S1:**
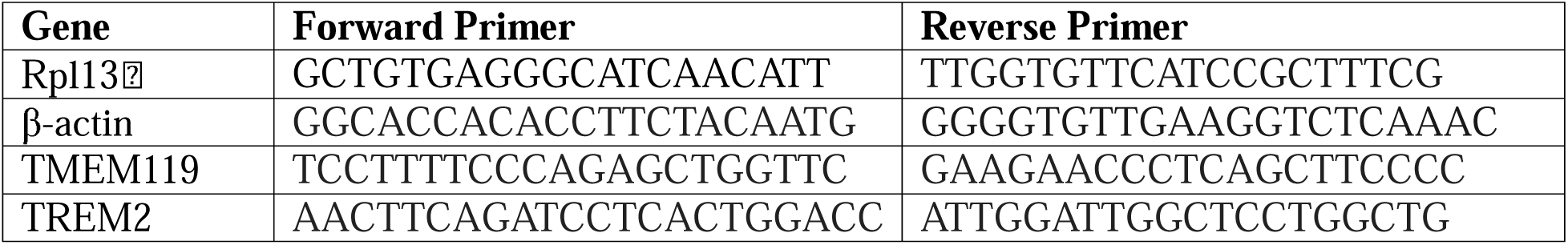
Primer sequences used for qPCR experiments.

**Figure S1:**
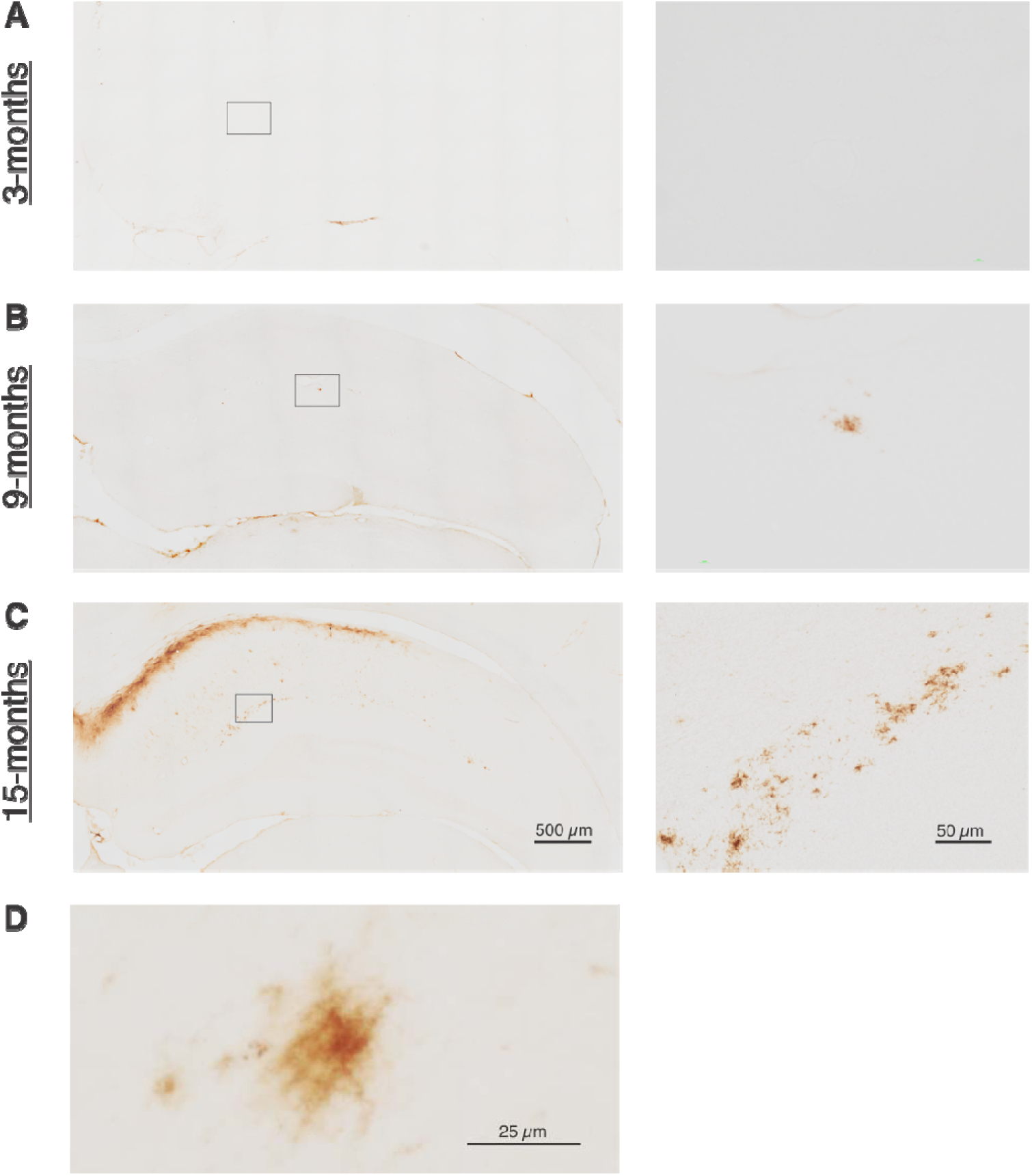
Immunohistochemistry of 82E1 for amyloid deposition in 3-, 9-, and 15-month APP/PS1 rats. A-C) Left panel images are 10 ⍰ stitches of the hippocampus (scale bar indicates 500 µm) and right panel images are 20 ⍰ captures of the outlined area (scale bar indicates 50 µm) in A) 3-month, B), 9-month, and C) 15-month APP/PS1 rats. D) Image taken at 40 ⍰ magnification depicting amyloid plaque (scale bar indicates 25 µm).

**Figure S2:**
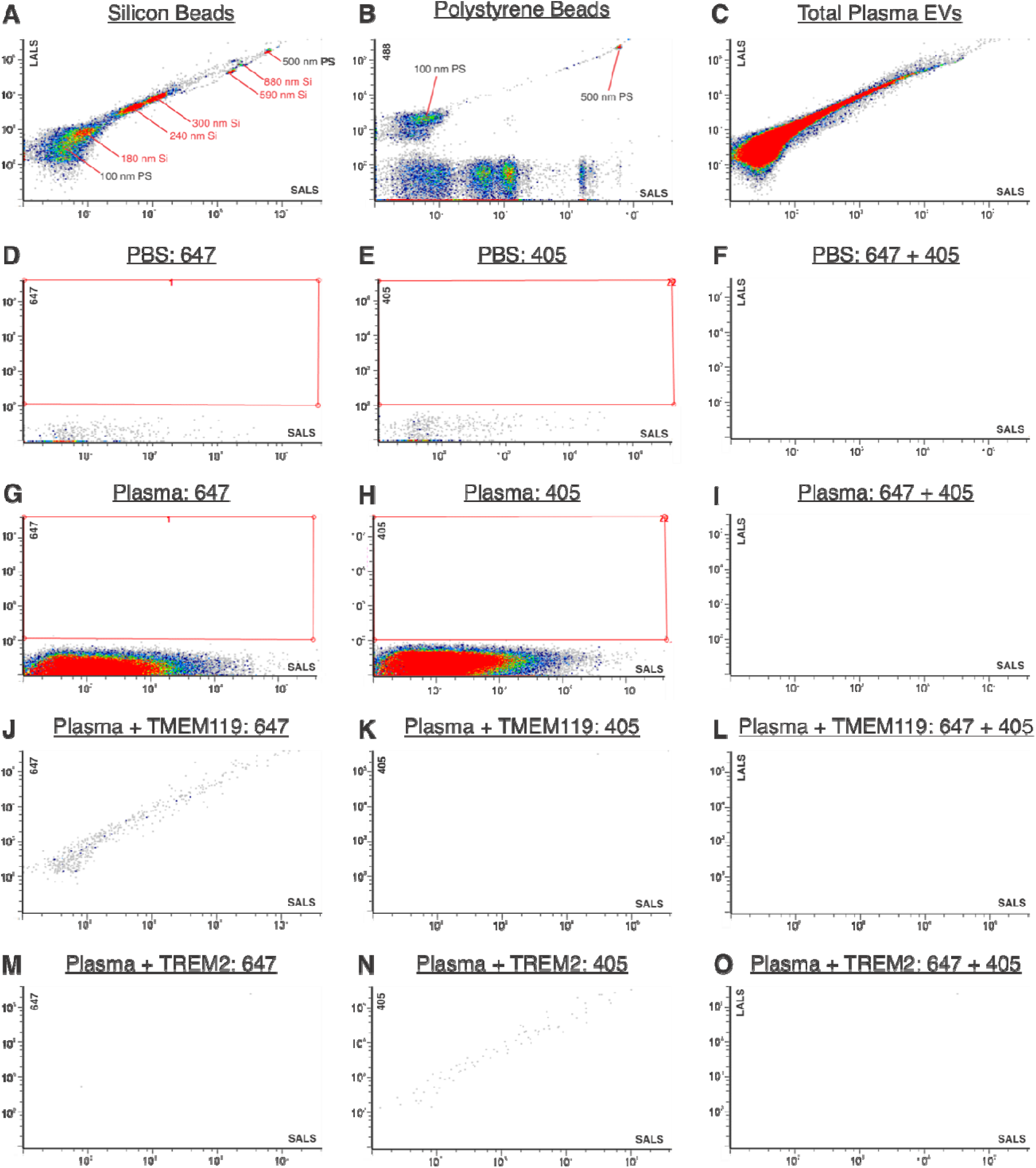
Nanoflow detection of TMEM119, TREM2, and standardized beads in rat plasma. A-B) Scatter plots of silicon beads (180 nm, 240 nm, 300 nm, 590 nm, and 880 nm) and 488-conjugated polystyrene beads (100 nm and 500 nm). C) Scatter plot of total plasma EVs. D-F) PBS in 647-, 405, and dual-gated channels. G-I) Plasma in 647-, 405-, and dual-gated channels. J-L) Plasma + TMEM119-647 antibody in 647-, 405-, and dual-gated channels. M-O) Plasma + TREM2-405 in 647-, 405-, and dual-gated channels. LALS, long angle light scatter; SALS, short angle light scatter.

**Figure S3:**
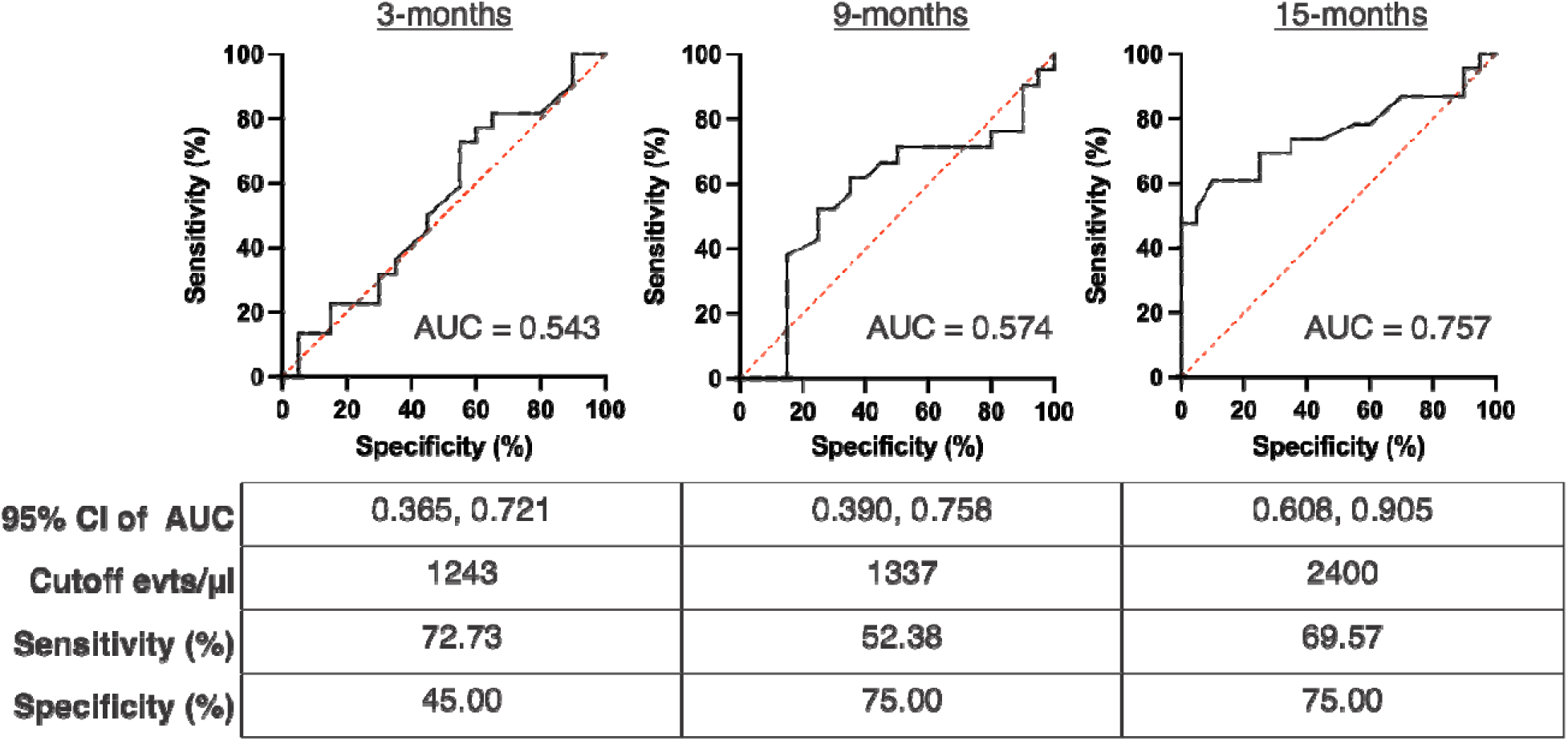
Discrimination of wildtype from APP/PS1 rats using TMEM119^+^/TREM2^+^ EVs. Receiver operating characteristic (ROC) curves of 3-, 9-, and 15-month rats and area under the curve (AUC) values.

**Figure S4:**
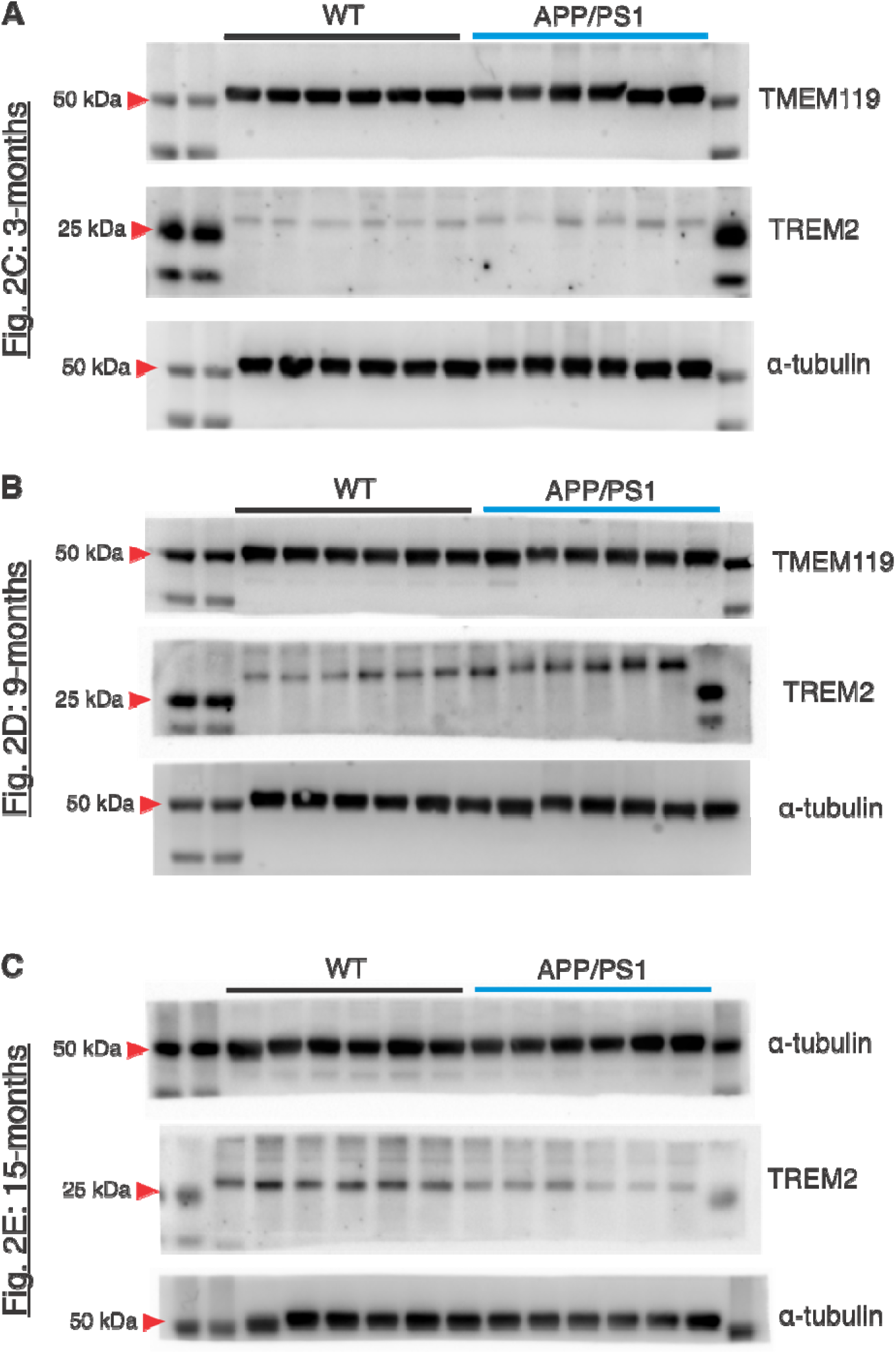
Hippocampus western blot imaging files.

**Figure S5:**
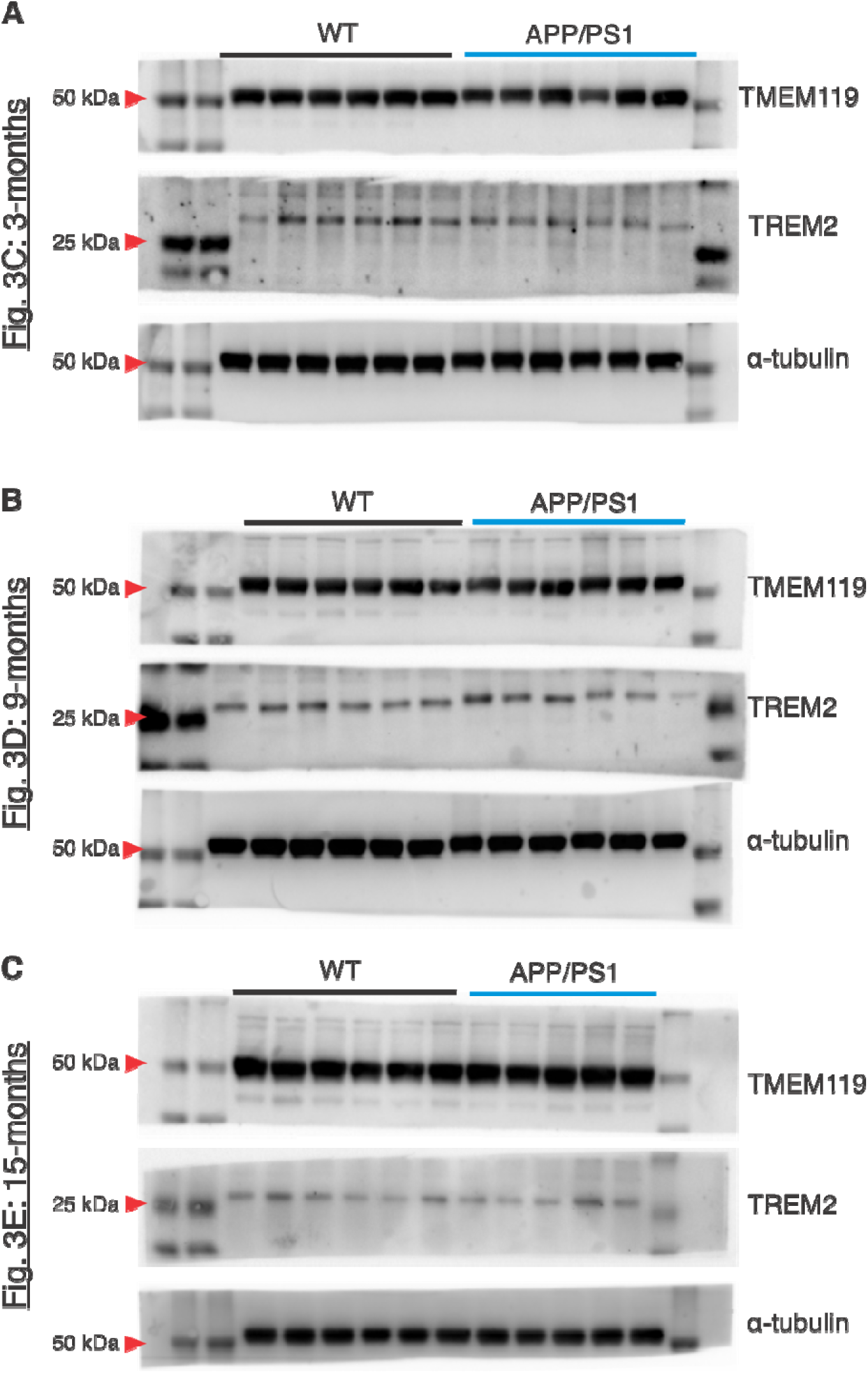
Corpus callosum western blot imaging files.

**Figure S6:**
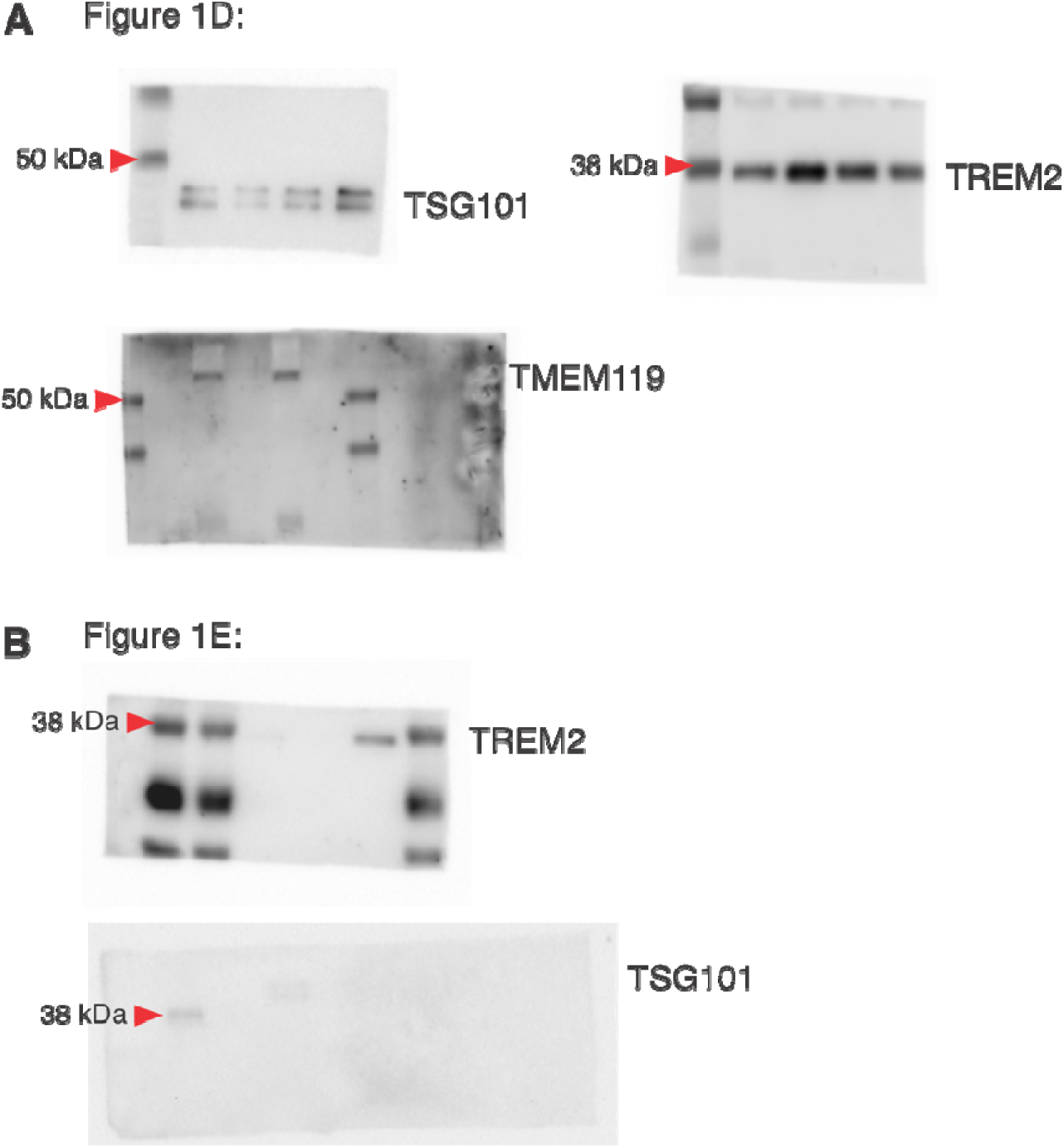
EV isolation western blot imaging files.

